# βPix sequesters IDOL and prevents LDL receptor degradation through a β_2_AR-regulated signaling pathway in Alport Syndrome

**DOI:** 10.1101/2020.11.06.372292

**Authors:** Ahmed Chahdi, Keyvan Yousefi, Jose Manuel Condor Capcha, Camila Iansen Irion, Guerline Lambert, Serene A. Shehadeh, Julian C. Dunkley, Yee-Shuan Lee, Aisha Khan, Melina Ramic, Nadja S. Andrade, Zane Zeier, Derek M. Dykxhoorn, Chryso Pefkaros Katsoufis, Michael Freundlich, Joshua M. Hare, Mary Nabity, Carolina Rivera, Anastasios Lymperopoulos, Lina A. Shehadeh

## Abstract

Alport syndrome (AS) is a rare disease of the glomerular basement membrane type IV collagen causing progressive renal failure. We reported increased accumulation of low-density lipoprotein (LDL) receptor (LDLR) and subsequent LDL cholesterol (LDL-C) uptake in renal tubular epithelial cells (TEC) in Alport mice, but the mechanisms regulating LDLR stability and function remain unknown. Here, we show that a selective β_2_-Adrenoceptor (β_2_AR) agonist, salbutamol, decreased LDLR levels and LDL-C uptake in Alport kidneys accompanied with reduced albuminuria and improved cardiac systolic and diastolic function. Similarly, salbutamol decreased LDL-C uptake in HK2 and HEK293 human renal epithelial cell lines, in smooth muscle cells from an X-linked hereditary nephropathy dog model (a large animal model of AS), and in TECs differentiated from AS patient-derived iPSCs. We show that the Rac1/Cdc42 guanine nucleotide exchange factor β1Pix blocked β_2_AR-induced LDLR degradation and, hence, increased LDL-C uptake. β1Pix also abrogated ubiquitination and degradation of LDLR induced by the inducible degrader of the LDLR (IDOL), an E3 ubiquitin ligase that promotes lysosomal LDLR ubiquitination and degradation. We identify a multimolecular complex comprised of βPix, IDOL, and LDLR and demonstrate that βPix counteracts β_2_AR-mediated LDLR degradation by sequestering IDOL. Our findings show βPix acts as a significant post-transcriptional regulator of IDOL-mediated LDLR degradation and identify β_2_AR activation as a potential treatment for Alport pathology.

---

Alport Syndrome (AS) is a hereditary nephropathy caused by mutations in genes encoding α-chains of type IV collagen *(Col4A)* leading to progressive glomerulonephritis and end-stage renal disease.^1^ In the absence of a cure, AS patients undergo dialysis and kidney transplant at early ages. ^2^ Therefore, there is a significant need to understand the mechanisms of renal failure in AS and identify new therapeutic approaches.

Abnormal cholesterol homeostasis results in excessive cholesterol and lipid accumulation in the kidney and is an established mechanism of renal injury. In recent work, we showed that low density lipoprotein receptor (LDLR) is highly expressed in the renal tubules of the *Col4a3^-/-^* Alport mouse and plays causative pathological roles.^3^ LDLR deletion was sufficient to extend the lifespan of the Alport mouse.^3^ Renal tubules of the Alport mouse were characterized by cholesterol accumulation, as well as dysmorphic mitochondria with defective bioenergetics. However, the exact mechanism(s) of this cholesterol nephrotoxicity and the relevant signaling effectors in human Alport pathology remain unknown.

The LDLR is a membrane-bound cell surface receptor complex with important endocytic functions for lipoproteins and a variety of extracellular ligands. ^4^ Owing to its central role in lipoprotein metabolism, LDLR levels are tightly regulated. Transcription of the LDLR is controlled by sterol regulatory element-binding proteins. Transcription rates increase when cellular cholesterol levels decline, to support increased endocytosis of cholesterol-rich LDL particles.^5, 6^ Ubiquitination of the LDLR is a major posttranscriptional determinant of receptor availability^7, 8^. The inducible degrader of the LDLR (IDOL) is an E3 ubiquitin ligase that promotes LDLR ubiquitination and subsequent lysosomal degradation of the receptor^9^. IDOL-mediated LDLR degradation preferentially targets the plasma membrane pool of receptors and is independent of the clathrin-mediated endocytosis of LDLR^8^. Given the high accumulation of LDLR in kidneys of Alport mice^3^ and the known cytotoxic effects of cholesterol in the kidney^10–14^, we sought to determine the molecular and signaling mechanisms that regulate LDLR stability in kidney cells *in vitro,* in renal tubular epithelial cells (TECs) from human cell lines and Alport patients iPSCs, and *in vivo* in Alport mice.

## RESULTS

### Acute stimulation of β_2_-Adrenoceptors by salbutamol reduces tubular LDL cholesterol (LDL-C) uptake and improves renal function in Alport mice

To evaluate the effects of acute β_2_AR stimulation on tubular LDL-C uptake and renal function, 7- week-old Alport mice were fasted overnight and subjected to a single intraperitoneal injection of 200μg salbutamol or equivalent vehicle control (schematic shown in Fig. 1a). Quantification of cAMP levels in kidney tissue 1h and 3h post salbutamol injection shows that single injection of salbutamol significantly elevated renal cAMP levels 1h post-injection, and dissipated within 3h post injection (Fig. 1b). This is consistent with the short duration of salbutamol activity. To evaluate the effects of acute β_2_AR activation on LDL-C influx *in vivo,* pHrodo red-labeled LDLC was injected via tail vein 1h after the salbutamol treatment (2h prior to sacrifice) and LDL-C uptake into the kidney tubules quantified by microscopic analysis. Our data showed that salbutamol markedly (~3-fold) reduced LDL-C influx (Fig. 1c-d, p<0.05) and KIM-1 expression (surrogate marker of kidney injury) (Fig. 1e-f, p<0.05) in renal tubules compared to control vehicle treatment. Moreover, urinary albumin-to-creatinine ratio (Alb/CRE) was significantly reduced in salbutamol-treated mice at 3h post-injection compared to the vehicle group and to baseline values (Fig. 1g), although plasma levels of blood urea nitrogen (BUN) and creatinine remained unchanged (Fig. 1h-i). Confirming these *in vivo* results, treatment of human kidney proximal tubular HK2 cells with 10 μM salbutamol showed a similar reduction in LDL-C influx *in vitro* (Fig. 1j). We further validated this finding in aortic smooth muscle cells from a dog with X-linked hereditary nephropathy (XLHN), a canine model of AS, where salbutamol at 10 μM significantly reduced LDL-C uptake (Fig. 1k).

**Figure 1.**
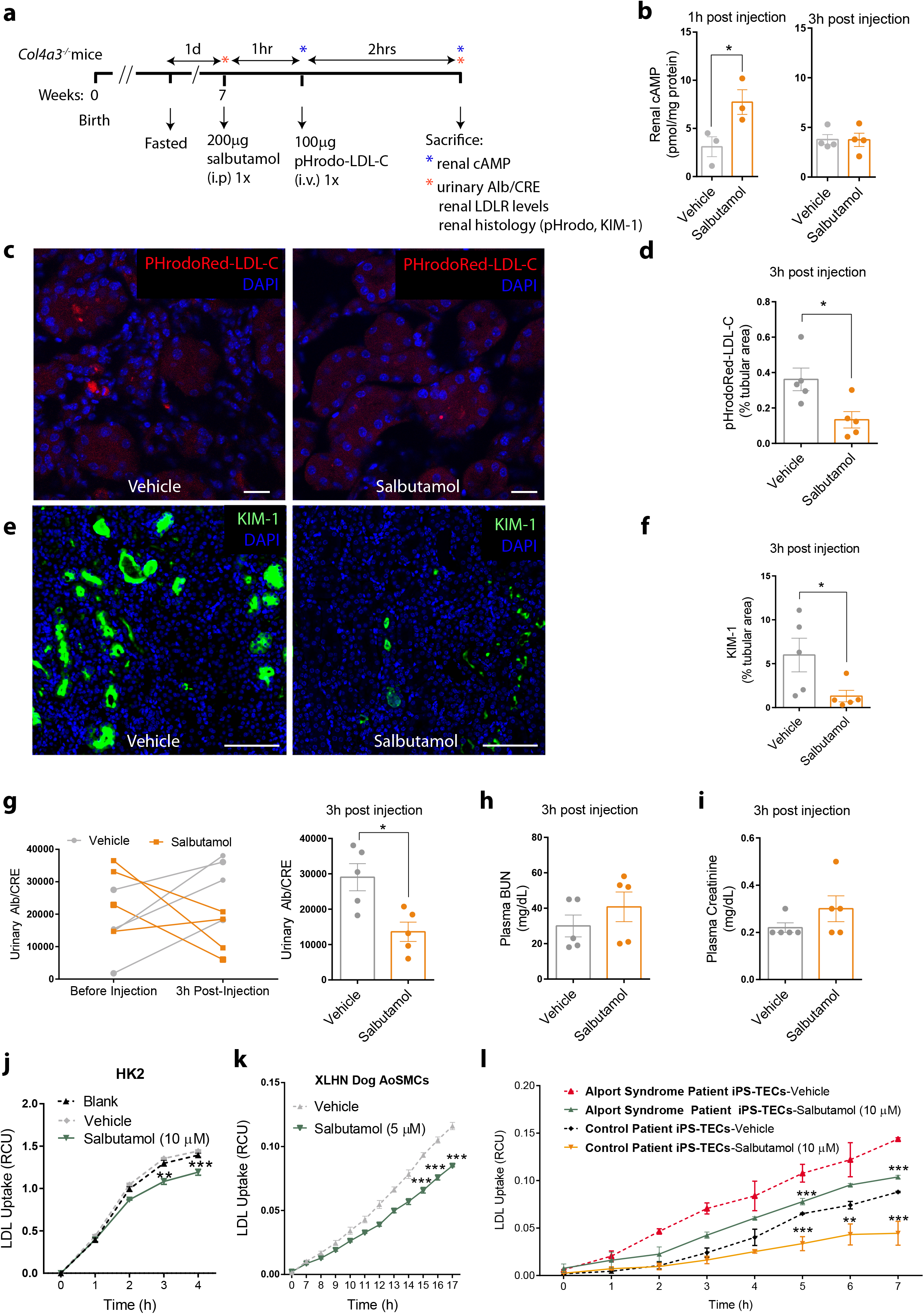
Short term treatment with salbutamol reduces LDL cholesterol influx and improves renal function in Alport mice. (a) Schematic of *in vivo* experiments. 7-week-old Alport mice were fasted overnight, injected intraperitoneally with a single bolus of 200μg salbutamol or DMSO as control vehicle and sacrificed 3 hours after treatment. Renal function, cAMP levels, and histology were evaluated at 1h and 3h post-injection. To evaluate the effects of acute β_2_AR stimulation on LDL cholesterol uptake *in vivo,* 100 μg pHrodo red-labeled LDL-C was injected via tail vein 1h after the treatment (2h prior to sacrifice). (b) Renal cAMP levels were significantly elevated at 1h but not 3h post-treatment. (c) Representative confocal images of kidney crosssections of treated mice and (d) the respective quantification of pHrodo red-labeled LDL-C show reduction in LDL-C influx in renal tubules of animals treated with salbutamol. Scale bar, 20μm. (e) Representative confocal images of kidney cross-sections of treated mice and (f) the respective quantification of Kidney Injury Molecule-1 (KIM-1) show reduction in LDL-C influx in renal tubules of animals treated with salbutamol. Scale bar, 100μm. (g) Urinary albumin-to-creatinine ratio (Alb/CRE) was significantly reduced in salbutamol-treated mice although plasma levels of (h) blood urea nitrogen (BUN) and (i) creatinine remained unchanged. Salbutamol treatment in (j) human kidney proximal tubular cells (HK-2), (k) aortic smooth muscle cells (AoSMCs) from a dog with X-linked hereditary nephropathy (XLHN), and (l) induced pluripotent stem cell-derived tubular epithelial cells (iPS-TECs) from a patient with Alport syndrome and a healthy individual significantly reduced LDL-C uptake as measured by live cell imaging. cAMP: Cyclic adenosine monophosphate; LDL: Low density lipoprotein; LDL-C: LDL Cholesterol. Data are mean±SEM. For *in vivo* experiments (b-i) N=3-5 mice/group were used. *p<0.05 and **p<0.01, using unpaired Student’s t-test. For *in vitro* experiments (j-l) N=3 wells/group was used and N=16 images/well were used for LDL uptake quantification. Each experiment was replicated at least three times. **p<0.01 and ***p<0.001, using Two-Way ANOVA.

### Differentiation of Alport patient-derived iPSCs into functional tubular epithelial cells (iPS-TECs)

We attempted to create an *in vitro* system with the highest similarity to the human Alport tubulopathy. To this end, we collected blood samples from four consenting patients (including one asymptomatic and three symptomatic Alport patients), 5 healthy individuals, and 3 individuals with non-Alport renal disease using established IRB protocols at the University of Miami. Demographic and diagnostic information of the subjects is shown in Online Table S1. We differentiated iPSCs from Alport patients, donor controls with non-Alport renal disease, and healthy volunteers into functional TECs as validated by flow cytometry and γ-Glutamyl Transferase (GGT) enzymatic assay, a marker of tubular epithelial cells^15, 16^. Online Fig S1 shows morphological changes in iPSCs prior to differentiation initiation compared to iPS-TECs on day 20 of differentiation in representative Alport and control donor cells. Representative validation results of iPSCs and iPS-TECs by flow cytometry are shown in Online Fig. S2 and by immunostaining in Online Fig. S3. We performed several differentiation experiments using control and Alport patient iPSCs. Prior to differentiation, 93-97% of the control or Alport iPSCs are triple positive for pluripotency markers TRA-1-60 (stem cell podocalyxin epitope), SOX2 (Sex Determining Region Y-Box 2), and SSEA4 (Stage-specific embryonic antigen-4), indicating valid iPSCs. However, 20 days post-differentiation, 60-70% of the derived cells are negative for both TRA-1-60 and SOX2, showing loss of pluripotency in the majority of the cells. Among the Tra-1- 60 negative/SOX2 negative cells, 56-76% stained positive for KSP1 (kidney-specific protein-1) and GGT1, indicating a very good yield of TECs. We performed flow cytometry analysis for iPSCs before and after TEC differentiation at least 6 times. We also validated the functionality of the generated TECs by GGT enzymatic assay as shown in Online Fig. S4 using HK2 cells, as a positive control cell line. According to our findings, iPSCs show minimal GGT activity prior to differentiation. However, GGT activity was significantly increased in iPS-TECs on day 20 of differentiation.

### Expression patterns of LDLR and KIM1 in Alport patient iPSCs

Flow cytometry was used to measure the expression levels of LDLR and kidney injury molecule (KIM1). We found that Alport patient iPSCs showed a mild increase (although statistically insignificant) in expression levels of LDLR compared to healthy control iPSCs (Online Fig. S5, p=0.2). Moreover, Alport-specific iPSCs displayed increased KIM1 expression relative to control donor cells (Online Fig. S6, p<0.05), indicating KIM1 as a tubular injury marker that can be detected in iPSCs and may represent a reliable marker for assessment of renal injury *in vitro.*

### Salbutamol reduces LDL cholesterol uptake in Alport patient iPS-TECs

We previously reported that cholesterol homeostasis was dysregulated in Alport mouse renal tubules.^3^ Therefore, we aimed to compare the baseline levels of LDL-C influx in AS patient specific-compared to donor control-derived iPS-TECs, and to evaluate the effects of β_2_AR stimulation by salbutamol. Our data showed that, consistent with the severity of Alport disease, cholesterol uptake is significantly higher in iPS-TECs from an Alport patient (subject #1), who had the most advanced renal pathology among the AS cohort (Fig. 1I). The level of LDL-C uptake was similar in cells from healthy donor controls and the AS patients with no renal insufficiency, indicating the potential association of kidney disease progression with the levels of LDL-C influx. The elevated levels of LDL-C influx in the iPS-TECs from this patient with end-stage renal disease is also in line with elevated LDL-C uptake seen in renal tubules of Alport mouse model on 129J background at 8 weeks of age which corresponds to advanced renal insufficiency. Validating our *in vivo* (in Alport kidneys) and *in vitro* (in HK2 cells and XLHN SMCs) findings about salbutamol regulation of LDL-C uptake, we show that stimulation of β_2_AR by salbutamol significantly decreased LDL-C uptake in Alport patient #1 iPS-TECs, as well as in the control donor cells as shown in Fig.1I.

### Chronic salbutamol treatment improved cardiac function in Alport mice

Echocardiographic analysis of cardiac function in Alport mice treated daily with salbutamol for 14 days, showed a significant improvement in systolic and diastolic cardiac function compared to the control vehicle group (schematic shown in Fig. 2a). Salbutamol treatment resulted in increased ejection fraction, stroke volume, cardiac output, and global longitudinal strain (GLS), reflecting a marked improvement in systolic function (Fig. 2b-e). Additionally, diastolic function was improved by salbutamol as evidenced by a significantly reduced isovolumetric relaxation time (IVRT, Fig. 2f), early mitral inflow velocity to mitral annular early diastolic velocity ratio (E/E’, Fig. 2h), and myocardial performance index (MPI, Fig. 2i). E/E’ ratio is a standard measure of LV filling pressures^17^ and higher MPI, which is inversely related to cardiac function, utilizes both systolic and diastolic time intervals to assess global cardiac dysfunction.^18^ There was no change in early to late ventricular filling velocities ratio (E/A, Fig. 2g). We found that 2 weeks of daily salbutamol injections significantly increased cAMP levels in the heart (p<0.01, Fig. 2j left panel) but not in the kidney tissue (Fig. 2j right panel). In line with the unchanged levels of cAMP in the kidney and despite the effects of acute treatment, we did not detect any significant improvements in the plasma or urinary measures of renal function (Fig. 2k-m) or *in vivo* tubular LDL uptake (Fig. 2n). Consistently, KIM-1 expression was not changed with 2-week salbutamol treatment (Online Fig. S7).

**Figure 2.**
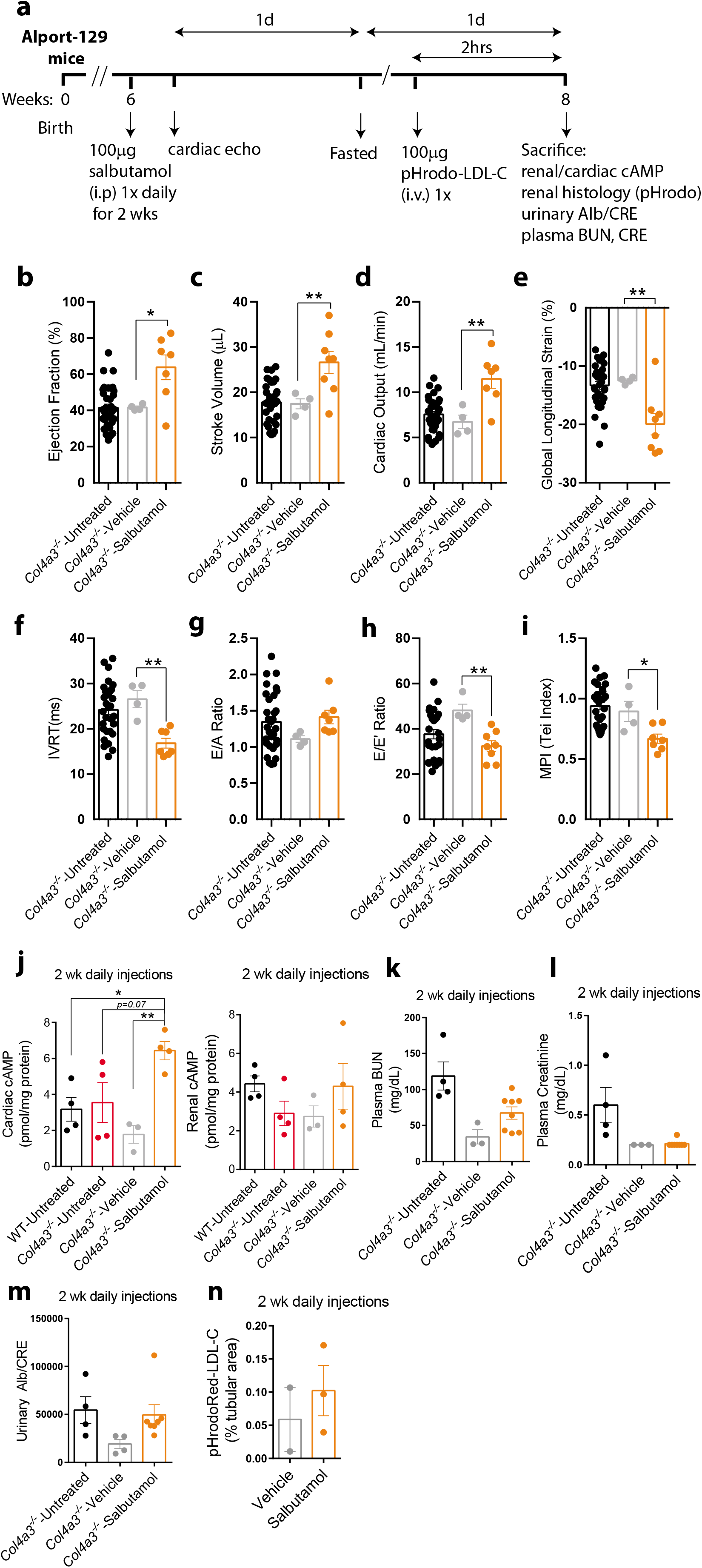
Chronic treatment with salbutamol improves cardiac but not renal function in Alport mice. (a) Schematic of *in vivo* experiments. Daily intraperitoneal injections of 100 μg/mouse salbutamol or equivalent DMSO as control vehicle were performed in 6-week-old Alport mice for 2 weeks. After the 2-week treatment, cardiac function, renal function, cAMP levels (renal and cardiac), and kidney histology were evaluated. To evaluate the effects of chronic β_2_AR activation on LDL cholesterol uptake *in vivo,* 100 μg pHrodo red-labeled LDL cholesterol (LDLC) was injected via tail vein 2h prior to sacrifice. Echocardiographic analysis of cardiac function shows that treating Alport mice with salbutamol significantly improved systolic (b-e) and diastolic (f-i) functional parameters compared to control vehicle-treated Alport mice. (j) Following the 2-week salbutamol treatment course, cAMP levels were significantly elevated in cardiac tissue (left panel) but renal cAMP remained unchanged (right panel). Chronic salbutamol treatment did not change measures of renal function (k-m) or tubular LDL uptake (n). Alb/CRE: Albumin-to-creatinine ratio; BUN: Blood urea nitrogen; cAMP: Cyclic adenosine monophosphate; IVRT: Isovolumic Relaxation Time; E/A: early to late ventricular filling velocities ratio; E/E’: early mitral inflow velocity to mitral annular early diastolic velocity ratio; LDL: Low density lipoprotein; LDL-C: LDL Cholesterol; MPI: Myocardial Performance Index or Tei Index. Data are mean±SEM. N=3-33 mice/group. *p<0.05 and **p<0.01 using One-Way ANOVA with Sidak’s post-hoc test.

### Salbutamol reduces LDLR levels

We previously observed increased levels of LDLR and increased LDL-C uptake in renal tubules in Alport mice.^3^ Therefore, herein we sought to determine the molecular mechanism(s) leading to the increase in LDLR protein levels. To determine the effect of the β_2_AR selective agonist, salbutamol, on LDLR protein levels *in vivo*, Alport mice were administered salbutamol (200 μg) or saline solution once and after 3 days, the kidney tissues were harvested and analyzed for LDLR expression by immunoblotting. The salbutamol-treated mice showed a marked decrease in LDLR protein levels relative to the control vehicle-treated mice (Fig. 3a). Similarly, stimulation of HEK293 cells expressing LDLR-GFP with salbutamol decreased LDLR protein levels after 3h stimulation (Fig. 3b), which is correlated with a decrease in LDL-C uptake (Fig. 3c). In HK2 cells, overexpression of the Rac1/Cdc42-GEF, β1Pix, increased LDL-C uptake and mitigated the inhibitory effect of salbutamol on LDL-C uptake, whereas the β1PixGEF-defective mutant, β1PixΔDH, showed no effect (Fig. 3c). Similar results were found using HepG2 cells (data not shown).

**Figure 3.**
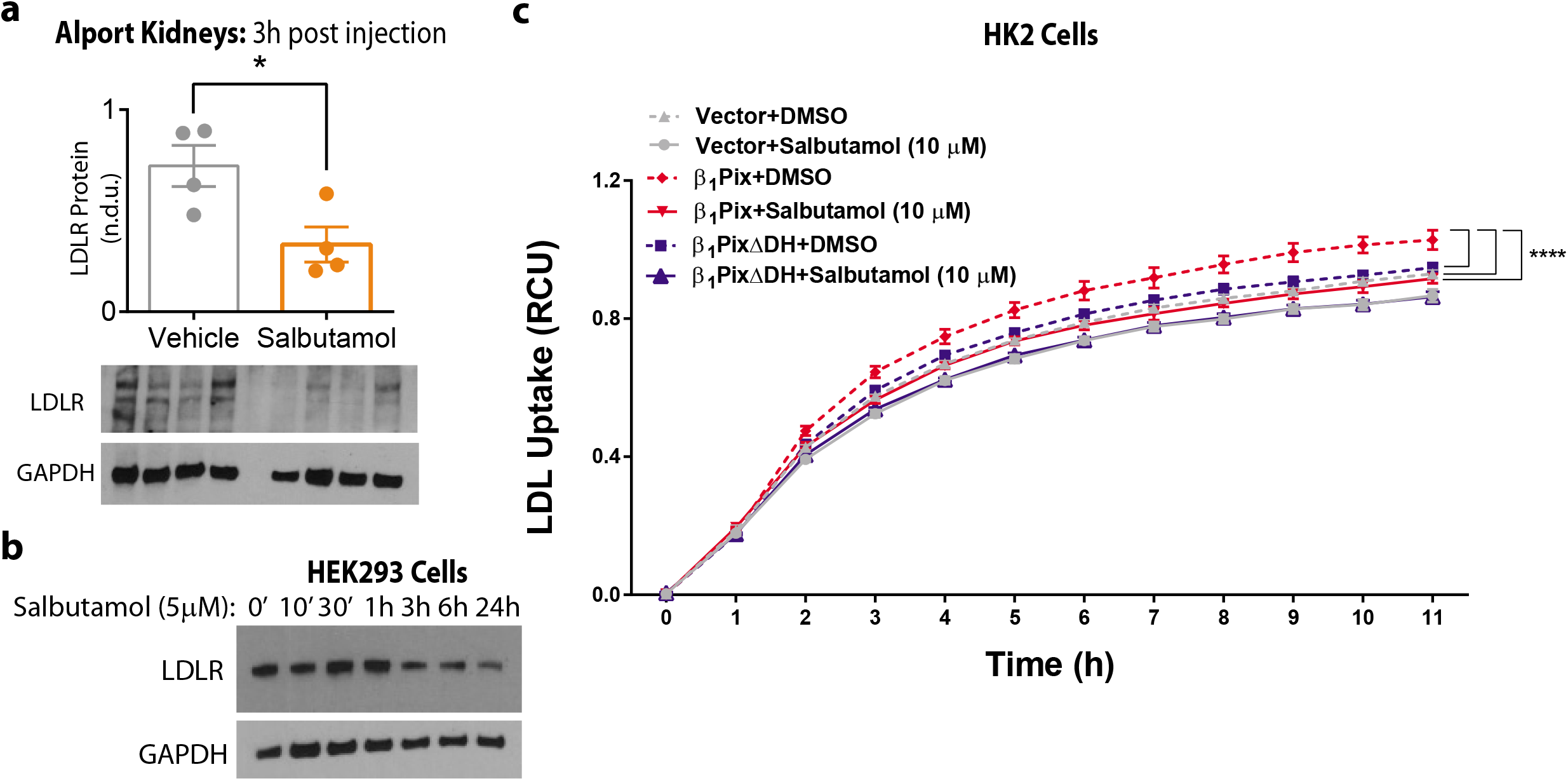
Salbutamol decreases LDLR levels. .(a) LDLR protein levels in Alport kidneys are reduced after 3 hours of 200μg Salbutamol injection. N=4 mice per group; *, p<0.05 by t-test. (b) HEK293 cells were stimulated with salbutamol (5 μM) for the indicated times. Total cell lysates were analyzed by immunoblotting as indicated. All Blots are representative of at least three independent experiments. (c) HK2 cells were transfected with empty vector, Mycβ_1_Pix or Mycβ_1_PixΔDH for 24h. Cells were incubated in serum-free media for 4h before stimulation with vehicle or salbutamol (10 μM) and LDL-C uptake was quantified. ****, p<0.0001 by 2-Way ANOVA with Dunnett multiple comparison test.

### βPix blocks the IDOL-mediated degradation of the LDLR

It has been shown that IDOL serves as E3 ubiquitin ligase that promotes ubiquitination and degradation of LDLR ^9^. Hence, this finding combined with our data described above raises the question of whether βPix modulates the IDOL-mediated degradation of LDLR. To examine this hypothesis, we used HEK293 cells because they have proven to be an excellent model to study LDLR ubiquitination and degradation^9^. The over-expression of HA-tagged ubiquitin in HEK293 cells expressing LDLR-GFP significantly increased the incorporation of HA-ubiquitin into LDLR compared to control cells (Fig. 4a). The ectopic expression of β1Pix led to a significant reduction in LDLR ubiquitination, whereas β1Pix mutant with defective GDP/GTP exchange activity yielded a very modest reduction in LDLR ubiquitination (Fig. 4a). Next, we tested the effect of β1Pix on IDOL-mediated LDLR degradation. HEK293 cells expressing LDLR-GFP were co-transfected with FLAG-IDOL and either Mycβ1Pix or Mycβ1PixΔDH. The expression of β1Pix protected LDLR from IDOL-mediated degradation, whereas the β1Pix-GEF defective mutant, β1PixΔDH, had only a minor effect (Fig. 4b). Furthermore, co-expression of β1Pix together with LDLR and IDOL inhibited degradation of the receptor in a dose-dependent manner (Fig. 4c).

**Figure 4.**
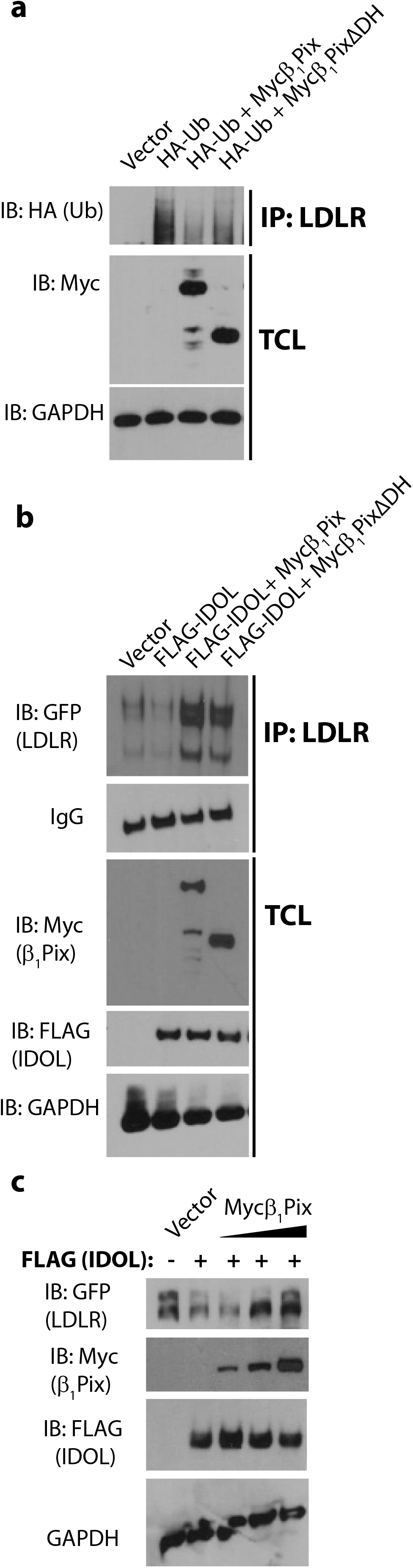
β1Pix inhibits IDOL-mediated ubiquitination and degradation of LDLR. (a) HEK293 cells expressing LDLR-GFP were transfected with empty vector or with an expression plasmid encoding HA-tagged ubiquitin alone or with the indicated Myc-tagged β1Pix constructs for 24h. The cell lysates were subjected to immunoprecipitation with anti-LDLR antibody. (b) HEK293 cells expressing LDLR-GFP were transfected with empty vector or with an expression plasmid encoding FLAG-tagged IDOL alone or with the indicated Myc-tagged β1Pix constructs. The cell lysates were subjected to immunoprecipitation with anti-GFP antibody. The expression levels of GFP-LDLR, FLAG-IDOL, and different forms of Myc-β1Pix in TCLs are shown. (c) HEK293 cells expressing LDLR-GFP were transfected with empty vector, expression plasmid for FLAG-IDOL, and increasing amounts of Myc-β1Pix. Total cell lysates were immunoblotted as indicated. The data are representative of at least three independent experiments.

### The βPix protein is a novel binding partner for LDLR

βPix was identified as a p21-activated kinase (Pak)-interacting exchange factor and was shown to be a GEF for Cdc42 and Rac1^19, 20^. Rho-GEFs activate Rho GTPases by catalyzing the exchange of GDP with GTP at the nucleotide binding site. In addition to Dbl homology (DH) and pleckstrin homology (PH) domains, βPix contains a Src homology 3 (SH3) domain responsible for binding Pak through a proline-rich region^20, 21^. βPix also has a leucine zipper domain for homodimerization^22^ and a GIT1 (G protein-coupled receptor kinase interactor 1) binding domain^21^.

To understand how βPix protect LDLR from IDOL-mediated degradation, we first sought to determine if βPix physically interacts with LDLR. We found that LDLR interacts with βPix. For example, Fig. 5a shows that Myc-tagged β1Pix was co-immunoprecipitated with LDLR from HEK293 cells. To gain further mechanistic insight into the physical interaction between β1Pix and LDLR, HEK293 cells were transiently transfected with additional mutated and truncated Myc-tagged β1Pix constructs. Cell lysates were immunoprecipitated with anti-LDLR antibody and resulting precipitates analyzed by immunoblotting with anti-Myc antibody. Deletion of PH or LZ domains did not affect the ability of β1Pix to bind to LDLR, whereas deletion of the DH domain significantly reduced its ability to be co-immunoprecipitated with LDLR (Fig. 5a) indicating that β1Pix dimerization is not required for β1Pix binding to LDLR, whereas the DH domain presence is necessary for β1Pix interaction with LDLR. To test whether β1Pix GEF activity within the DH domain is required for β1Pix interaction with LDLR, we transfected HEK293 cells with Myc-β1Pix or Myc-β1PixDHm(L238R, L239S) mutant unable to catalyze GDP/GTP exchange on Rac ^20^. Fig. 5b, shows that the double point mutation within the DH domain, β1PixDHm(L238R, L239S), strongly reduced the ability of β1Pix to bind to LDLR, indicating that β1Pix/LDLR association is dependent on β1Pix GEF activity. The SH3 domain is essential for βPix binding to Pak1 ^21^. Thus, our finding that the association of β1PixSH3m(W43K) with LDLR was indistinguishable from that of wild type β1Pix indicates that Pak1 binding is not essential for β1Pix association with LDLR (Fig. 5b).

**Figure 5.**
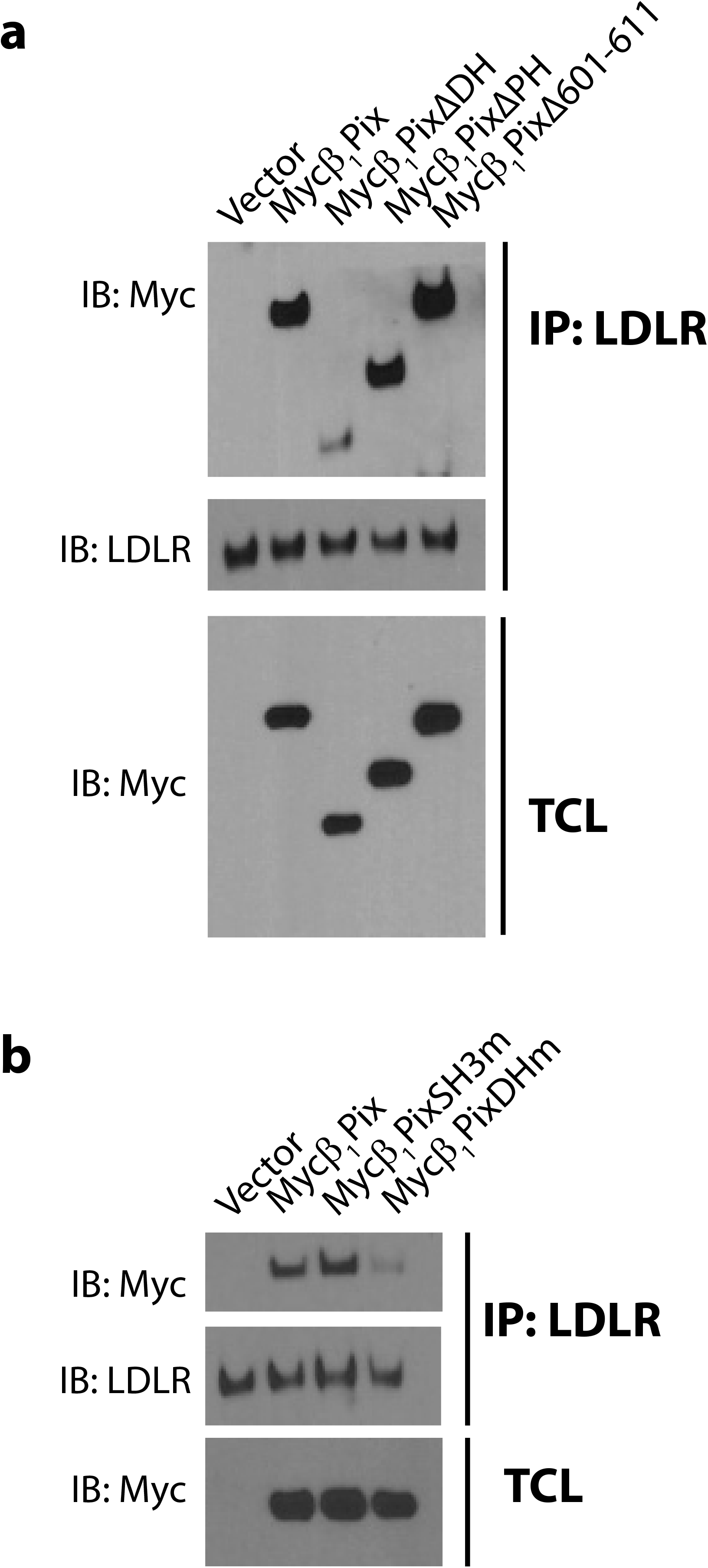
β1Pix GEF activity is required for its interaction with LDLR. (a) HEK293 cells were transfected with empty vector, β1Pix, β1Pix ΔDH, β1Pix ΔPH or β1Pix Δ(602-611), or with (b) empty vector, β1Pix, β1PixSH3m or β1PixDHm for 24 h. Total cell lysates were subjected to immunoprecipitation (IP) using anti-LDLR antibody and immunoblotting (IB) as indicated. The data are representative of at least three independent experiments.

### βPix rescues LDLR from degradation through IDOL sequestration

To determine whether β_2_AR modulates βPix/LDLR interaction at normal physiological levels, HEK293 cells were stimulated by salbutamol at different time points. Endogenous LDLR was immunoprecipitated from cell lysate using anti-LDLR antibody and the immunocomplexes were analyzed by Western blotting for βPix (Fig. 6a). Salbutamol induced a very rapid increase (2 minutes) in βPix association with LDLR, and this association reached its peak at 10 minutes post treatment and declined over the subsequent 30 minutes. Activation of β_2_AR elicits cAMPdependent signaling, therefore we tested the effect of adenylate cyclase on LDLR/βPix association. Fig. 6b shows that direct activation of adenylate cyclase by forskolin results in a rapid increase in endogenous LDLR/βPix association followed by a decrease after 30 minutes stimulation comparable to the kinetic of LDLR/βPix association induced by salbutamol.

**Figure 6.**
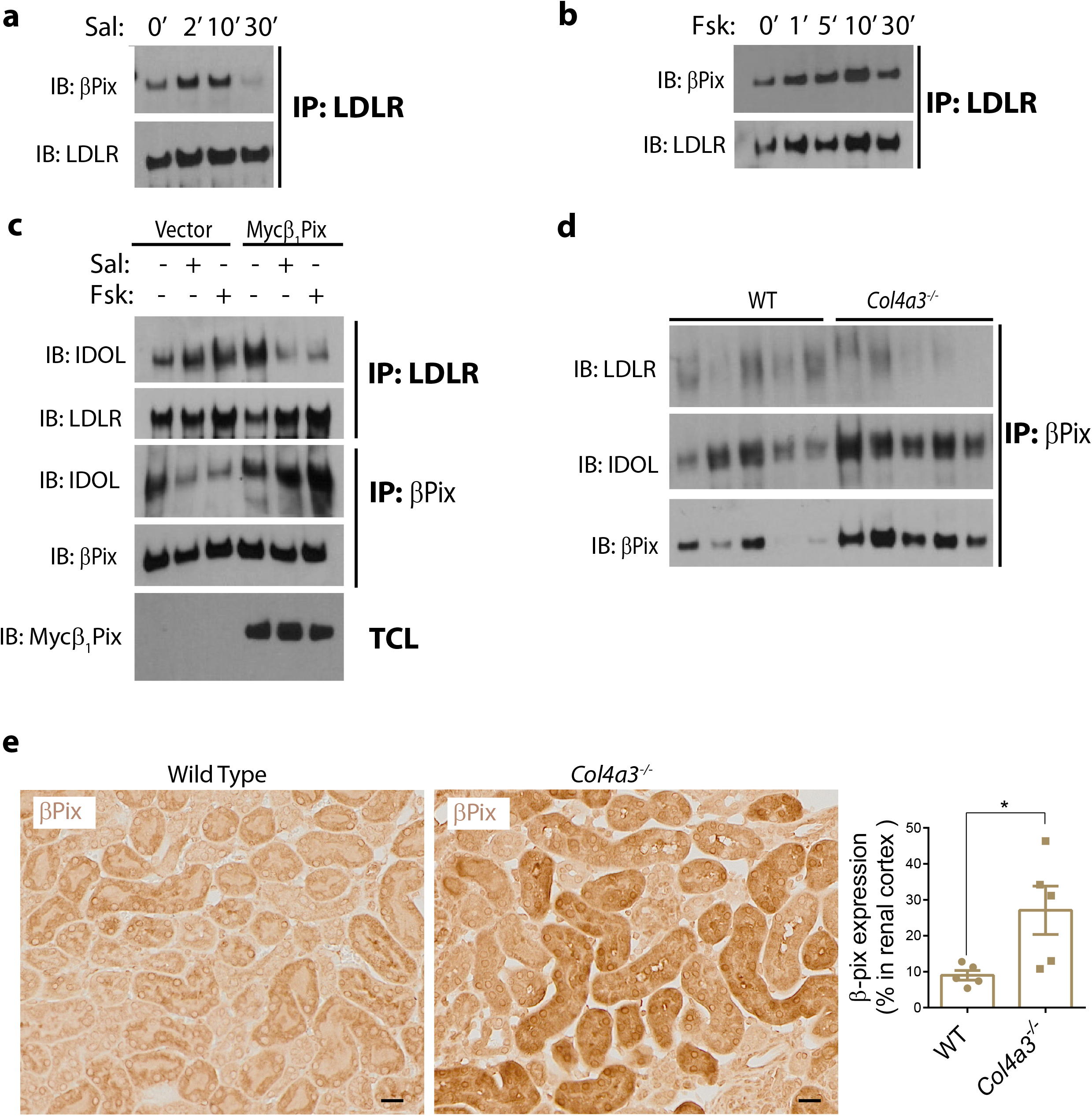
β1Pix rescues LDLR from degradation through IDOL sequestration. HepG2 cells were treated with IBMX (100 μM) before stimulation with 5 μM of salbutamol (a) or forskolin (b) for the indicated times. The cell lysates were subjected to immunoprecipitation using anti-LDLR antibody followed by immunoblotting with antibodies against βPix and LDLR. (c) HepG2 cells were transfected with empty vector or Myc-β1Pix for 24h. Cells were treated with IBMX (100 μM) for 30 minutes before stimulation with 5 μM of salbutamol or forskolin for 10 minutes. The cell lysates were subjected to immunoprecipitation using antibodies against LDLR or βPix followed by immunoblotting as indicated. (d) kidney tissues from WT and Alport mice were homogenized and subjected to immunoprecipitation using anti-βPix antibody followed by immunoblotting with antibodies against LDLR, βPix and IDOL. (e) βPix expression is increased in renal tubules of Alport mice relative to control mice as shown by immunostaining. N=5 mice per group; *, p<0.05 by t-test. Data shown are representative of at least three independent experiments.

To explore the underlying mechanism by which βPix protects LDLR from IDOL-mediated degradation, we examined the interaction dynamics between βPix, LDLR, and IDOL in HepG2 cells after β_2_AR stimulation. Functionally, β_2_AR preferentially couples to Gs and stimulates cAMP formation through activation of adenylate cyclase^23^. We found that stimulation of the cells with salbutamol or direct stimulation of adenylate cyclase by forskolin results in a strong increase in IDOL binding to LDLR and a simultaneous decrease of IDOL binding to β1Pix (Fig. 6c). Furthermore, in cells ectopically expressing β1Pix, both salbutamol and forskolin induced IDOL release from LDLR and increased its binding to β1Pix (Fig. 6c), indicating that salbutamol induces β1Pix-mediated sequestration of IDOL hence protecting LDLR from degradation. To support our hypothesis, we performed co-IP experiment on kidney tissues obtained from WT and Alport mice. Results show that in Alport mice IDOL association with βPix is increased while LDLR binding to βPix is decreased compared to WT mice (Fig. 6d). Furthermore, we found that Alport mice express more βPix compared to WT mice as shown by immunoprecipitation (Fig. 6d, bottom panel) and immunostaining (Fig. 6e).

## DISCUSSION

Nephroprotection through activation of β_2_AR signaling is lately emerging as a promising field of investigation.^24^ β_2_ receptors are expressed in multiple parts of the kidney including proximal and distal convoluted tubules, glomeruli, and podocytes.^24^ β_2_AR signaling has been extensively investigated in lung, cardiovascular, and muscular diseases^25–27^, but its potential roles in renal pathophysiology remain understudied. Specifically, β_2_AR signaling has never been investigated in AS.

Formoterol, a long-acting β_2_ selective agonist, was shown to improve the recovery of renal function after acute kidney injury (AKI) by activating proximal tubule β_2_AR and inducing mitochondrial biogenesis.^28, 29^ A recent study reported the prognostic significance of renal β_2_AR expression in clear cell renal cell carcinoma where reduced β_2_AR levels were associated with poor prognosis.^30^

Excessive lipid accumulation in renal cells is an established mechanism of renal injury. In *Col4a3^-/-^* mouse model of AS, we have reported increased expression of renal tubular osteopontin (OPN) and LDLR, and enhanced lipid accumulation in renal tubules.^3^ In addition, we have shown benefits for downregulating OPN and LDLR in this model^3^. OPN inhibition has been previously linked to prolonged cAMP generation under β_2_AR stimulation in an osteoblastic cell line MC3T3-E1.^31^ We also recently reported that OPN deficiency can enhance β_2_AR-mediated cAMP levels and anti-fibrotic gene program in cardiac cells.^32^

In the present study, we show that β_2_AR agonism using salbutamol reduces LDL cholesterol uptake *in vitro* in HK2 and iPS-TECs from Alport patients as well as canine aortic smooth muscle cells isolated from dogs with X-linked Hereditary Nephropathy (XLHN), the equivalent of human AS^33,34^ Similarly, short term stimulation of β_2_AR reduces LDL cholesterol uptake in Alport mouse renal tubules. Moreover, we found significant improvements in cardiac and renal measures following salbutamol therapy. These data suggest that β_2_AR agonism may be a viable strategy to prevent lipid toxicity in renal diseases associated with lipid accumulation, including AS.^35^

Mechanistically, we describe a novel relationship between β_2_AR signaling, βPix, and LDLR. The ability of β_2_AR-induced LDLR degradation is markedly reduced in cells that express βPix. Following endocytosis, LDLR is targeted for lysosomal degradation via IDOL an E3 ubiquitin ligase that promotes LDLR ubiquitination and degradation.^9^ To investigate whether activation of β_2_AR affects LDLR expression *in vivo,* we treated Alport mice with salbutamol. 3h after injection, salbutamol reduced LDLR protein levels in the kidneys, confirming our *in vitro* data. In view of our data showing that ectopic expression of β1Pix reversed salbutamol-induced LDLR degradation, we reasoned that βPix counteracts IDOL-mediated ubiquitination and subsequent LDLR degradation by preventing the binding of IDOL to LDLR. Overexpression of β1Pix not only blocked IDOL-mediated degradation of LDLR but increased LDLR stability (compare lane#1 and lane#3, Fig. 3b), suggesting that βPix stabilizes endogenous LDLR. A β1Pix mutant that is unable to bind to LDLR is less effective in blocking salbutamol-induced LDLR degradation and decreased cholesterol uptake. The SH3 domain mediates binding of βPix to a polyproline stretch in Pak1 ^19, 20^. Thus, our finding that SH3m(W43K) mutation did not block LDLR binding to β1Pix indicates that Pak1 binding to β1Pix is not essential for the β1Pix/LDLR association. However, the β1Pix GEF defective mutant, β1PixDHm(L238R, L239S), that is unable to catalyze GDP/GTP exchange on Rac ^20^ was unable to bind to LDLR indicating that LDLR binding to LDLR requires βPix GEF activity.

Stimulation of β_2_AR by salbutamol or forskolin increased IDOL binding to LDLR and decreased binding of IDOL to βPix indicating that βPix rescues LDLR from degradation by sequestering IDOL. This molecular mechanism explains the increased stability of LDLR in Alport mice compared to WT mice. The protective effect of βPix is amplified by its increased expression in Alport mice.

In conclusion, our study discovered what we believe is a new pathway through which βPix promotes LDLR stability. It does so by sequestering IDOL hence preventing LDLR from degradation. This finding represents a novel molecular mechanism explaining the excessive amounts of LDLR present in Alport mice^3^. This novel β_2_AR signaling pathway modulates the interaction between LDLR/βPix/IDOL, and hence cholesterol uptake in renal tubules. Based on the importance of lipid toxicity on renal pathology, modification of the LDLR via the β_2_AR couldbe a safe therapeutic approach to treat chronic kidney disease and related cardiac dysfunction in individuals with AS and potentially other renal diseases.

## Statistical analysis

For all experiments, N refers to the number of individual mice or individual culture plates. All data are expressed as mean ± S.E.M. P-values were calculated using unpaired Student’s t-test when comparing two groups. For data analysis of more than two group comparisons, p-values were calculated using ANOVA and corrected for multiple comparisons. Two-Way ANOVA is used for time course studies (LDL-C influx experiment). Repeated symbols represent p-values of different orders of magnitude, i.e. *p < 0.05, **p < 0.01, ***p < 0.001, ****p < 0.0001.

## Acknowledgments

This work was supported by the following grants to LAS: National Institute of Health (1R01HL140468) and the Miami Heart Research Institute. KY is a recipient of AHA predoctoral fellowship (18PRE33960070). JCD is a recipient of NIH Diversity Supplement Award (R01HL140468-02S1). This work was supported by NIH grant 1S10OD023579-01 for the VS120 Slide Scanner housed at the University of Miami, Miller School of Medicine Analytical Imaging Core Facility.

---

**Supplemental Figure S1.**
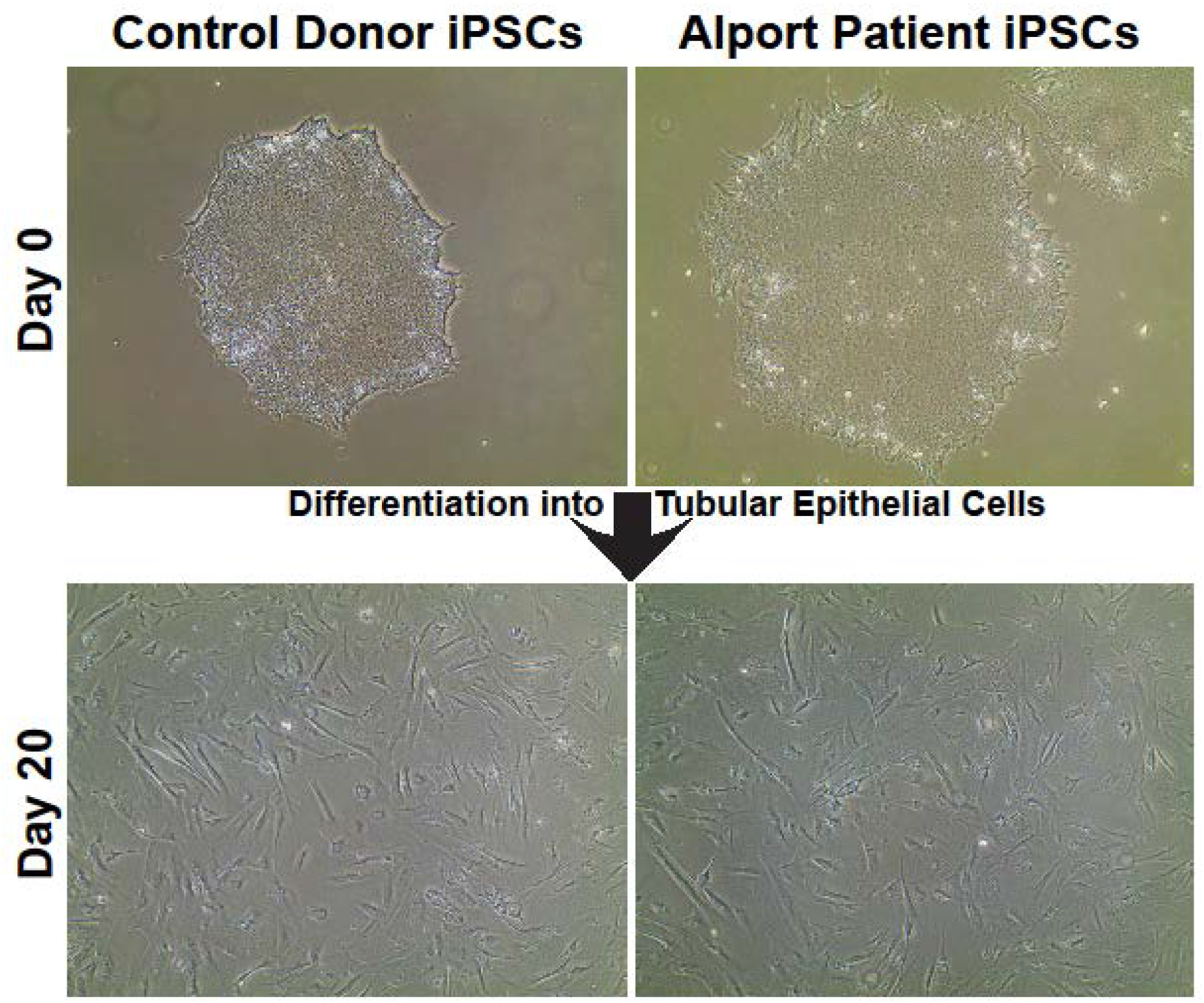
Morphology of iPSCs and iPS-TECs. Representative images of control donor (left panels) and Alport patient (right panel) iPSCs are shown prior to differentiation (top panels) and on day twenty following differentiation (lower panels). The morphological changes support differentiation of iPSCs into polarized TECs. iPS-TECs: induced pluripotent stem cell-derived tubular epithelial cells.

**Supplemental Figure S2.**
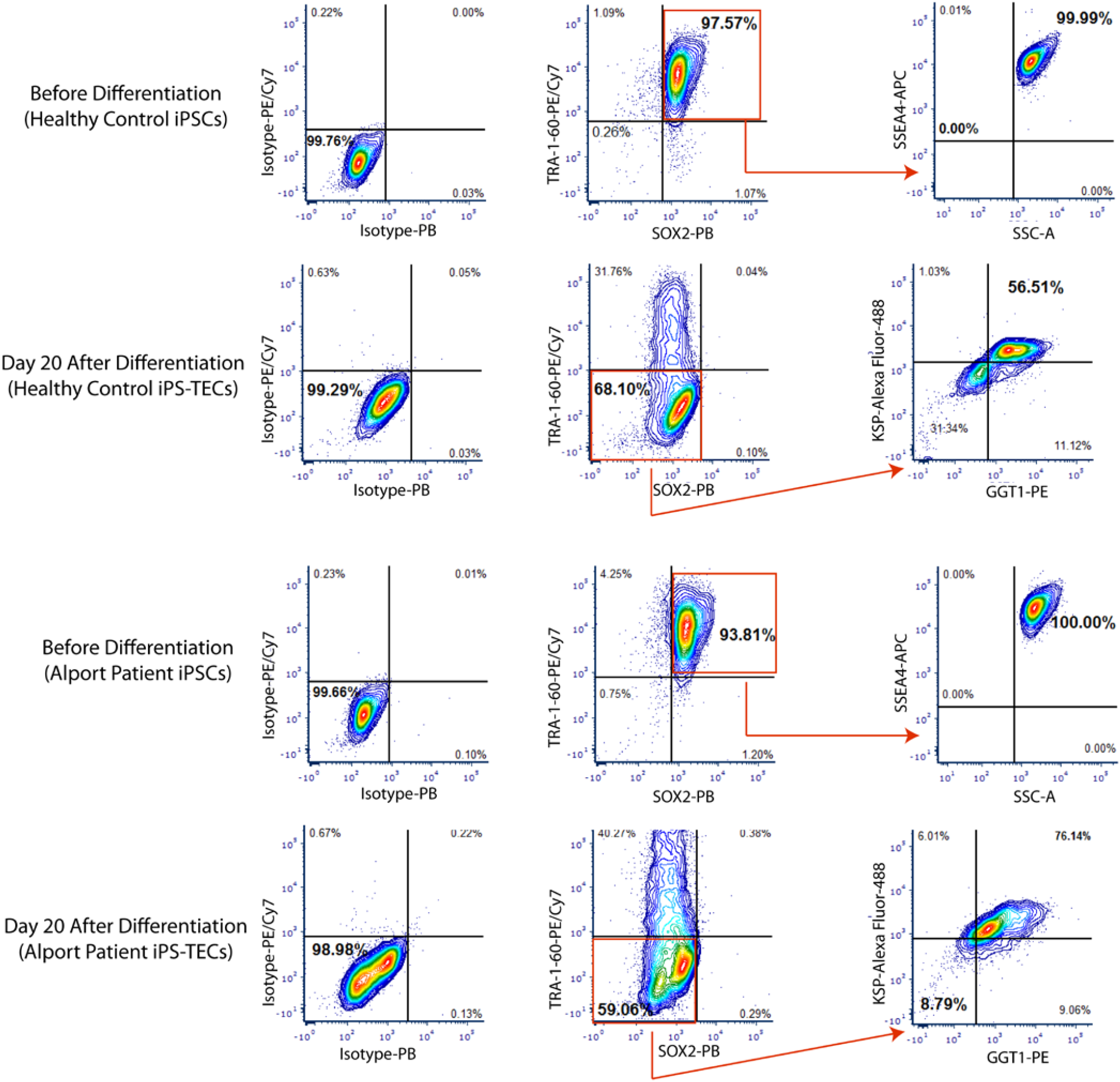
Successful reprogramming of PBMCs into iPSCs and differentiation into TECs. PBMCs from healthy controls and Alport patients were isolated and re-programmed into iPSCs as shown by expression of the pluripotency markers TRA-1-60 (cell surface), SOX2 (nuclear), and SSEA4 (cell surface). After differentiation into TECs, the pluripotency markers are lost and the cells express the renal tubular marker KSP1 and GGT. PBMCs: peripheral blood mononuclear cell; iPSCs: induced pluripotent stem cells; TECs: tubular epithelial cells; KSP1: kidney-specific protein-1; GGT: Gamma Glutamyl Transferase.

**Supplemental Figure S3.**
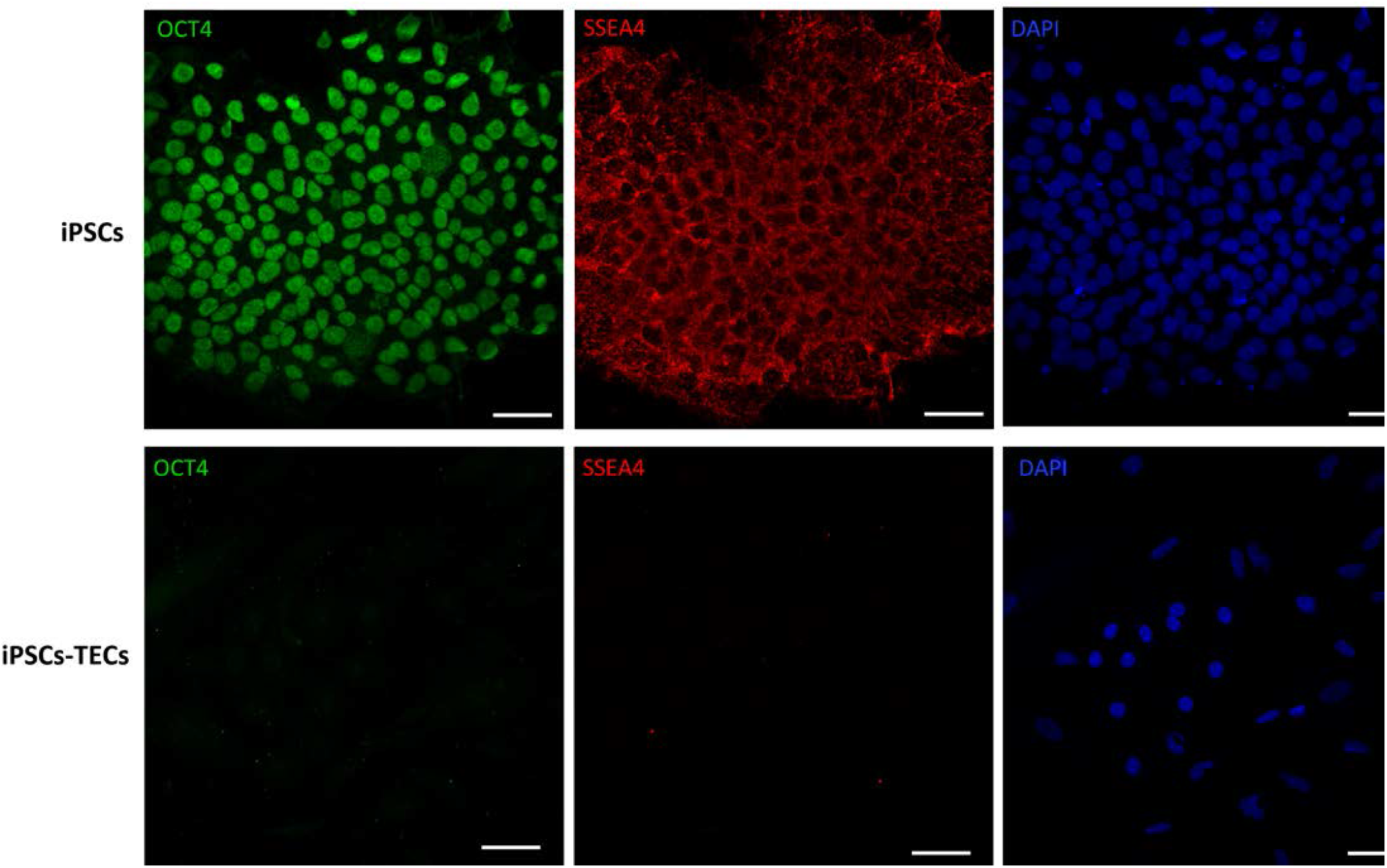
Successful reprogramming of PBMCs into iPSCs. Control donor PBMCs were isolated and re-programmed into iPSCs as shown by expression of the pluripotency markers SSEA (cytoplasmic) and OCT4 (nuclear). After TEC differentiation, the pluripotency markers are lost. Shown are representative confocal images. Scale bar, 20 μm.

**Supplemental Figure S4.**
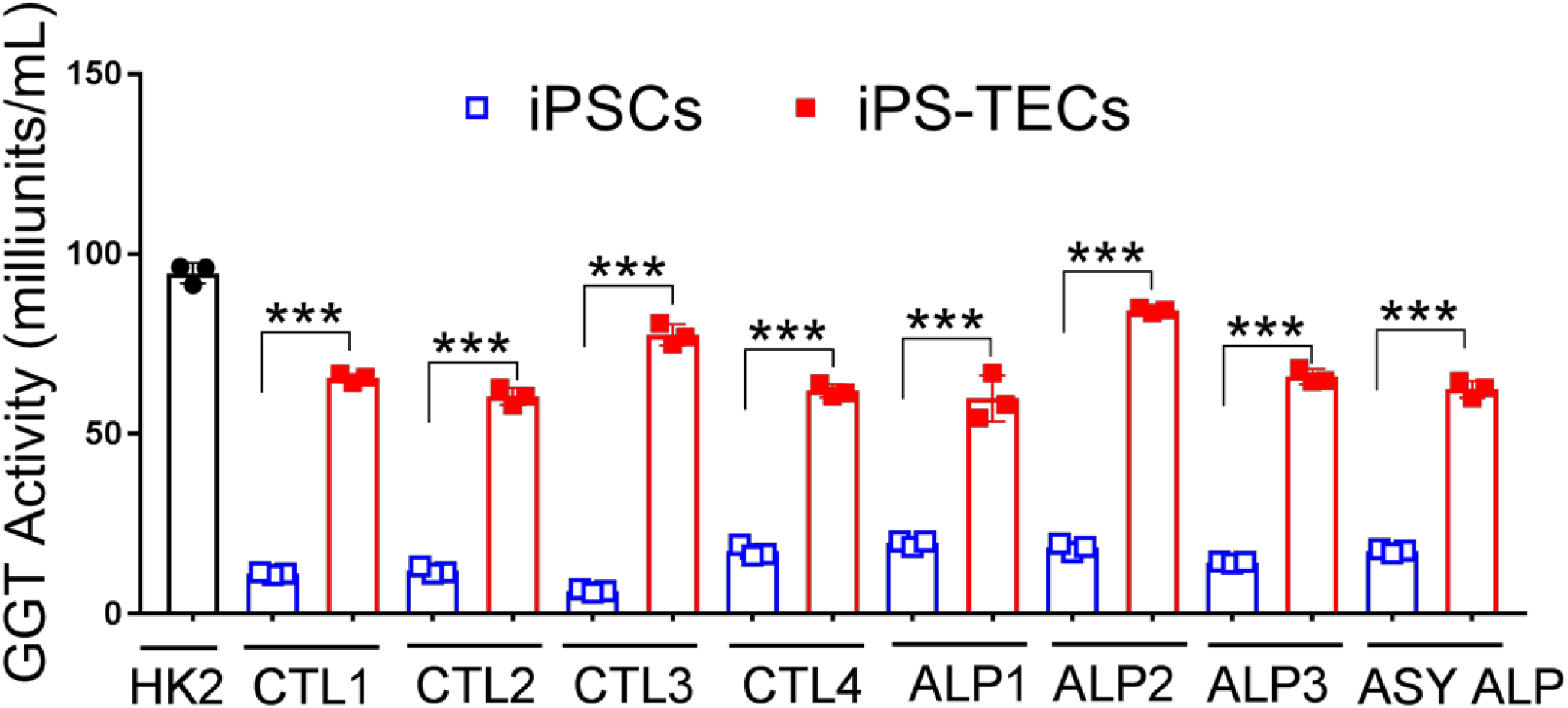
Alport patient and control donor iPS-TECs gain γ-glutamyl transferase (GGT) activity. GGT enzymatic activity, as a marker of tubular epithelial cells, is lacking in control donor or Alport patient iPSCs prior to differentiation. However, on day 20 of differentiation, the derived iPS-TECs show significantly increased GGT activity, validating the TEC commitment of the derived cells. Data are mean±SEM. The experiment was replicated at least three times. CTL: Control; ALP: Alport; ASY: Asymptomatic; ***p<0.001 using Unpaired Student’s t-test.

**Supplemental Figure S5.**
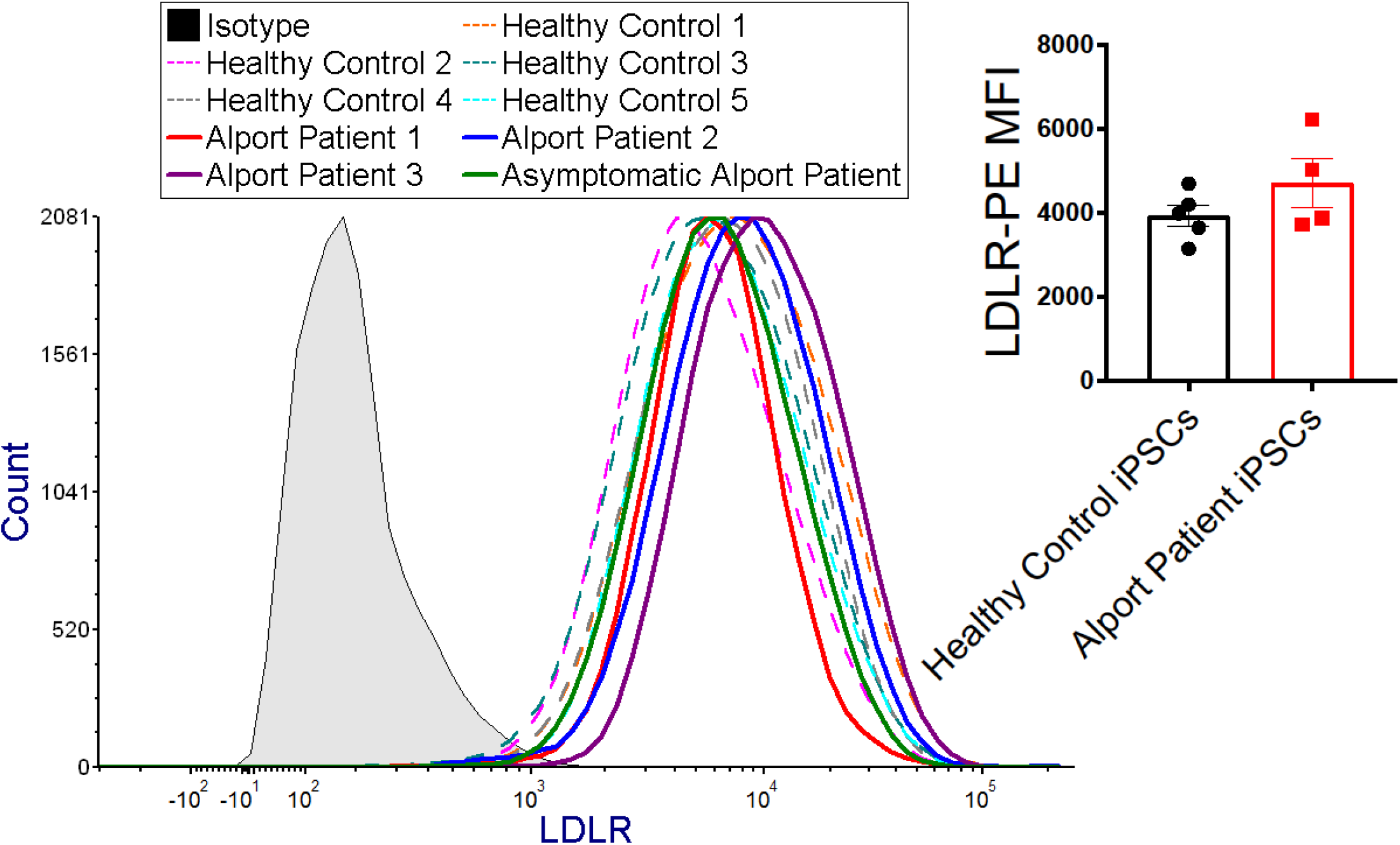
LDLR expression in Alport patient iPSCs. Flow cytometry findings showed that Alport patient and healthy control iPSCs express similar levels of LDLR. Data are mean±SEM. At least 10000 events per sample are analyzed and the final analysis is performed gating pluripotent (SOX2^+^TRA-1-60^+^) cell population. N=4 Alport patients and N=5 healthy control subjects. The experiment is replicated at least two times. AS: Alport Syndrome; MFI: Mean Fluorescent Intensity. No statistical significance was detected using Unpaired Student’s t-test.

**Supplemental Figure S6.**
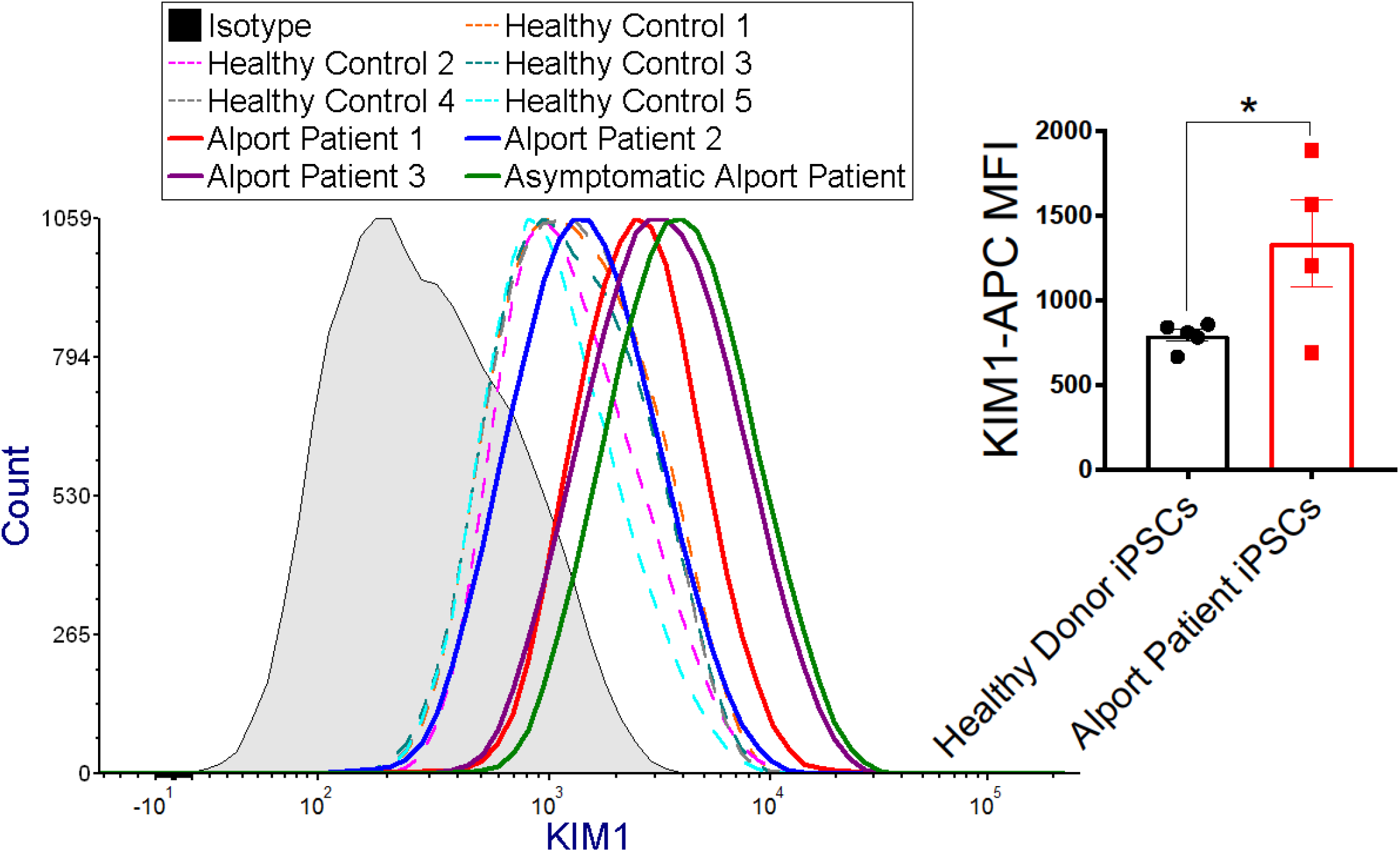
Alport patient iPSCs overexpress KIM1. Flow cytometry analysis showed that Alport patient iPSCs express higher levels of kidney injury molecule (KIM1) compared to healthy control donor cells. Data are mean±SEM. At least 10000 events per sample are analyzed and the final analysis is performed gating pluripotent (SOX2^+^TRA-1-60^+^) cell population. N=4 Alport patients and N=5 healthy control subjects. The experiment was replicated at least two times. AS: Alport Syndrome; MFI: Mean Fluorescent Intensity. *p< 0.05 using Unpaired Student’s t-test.

**Supplemental Figure S7.**
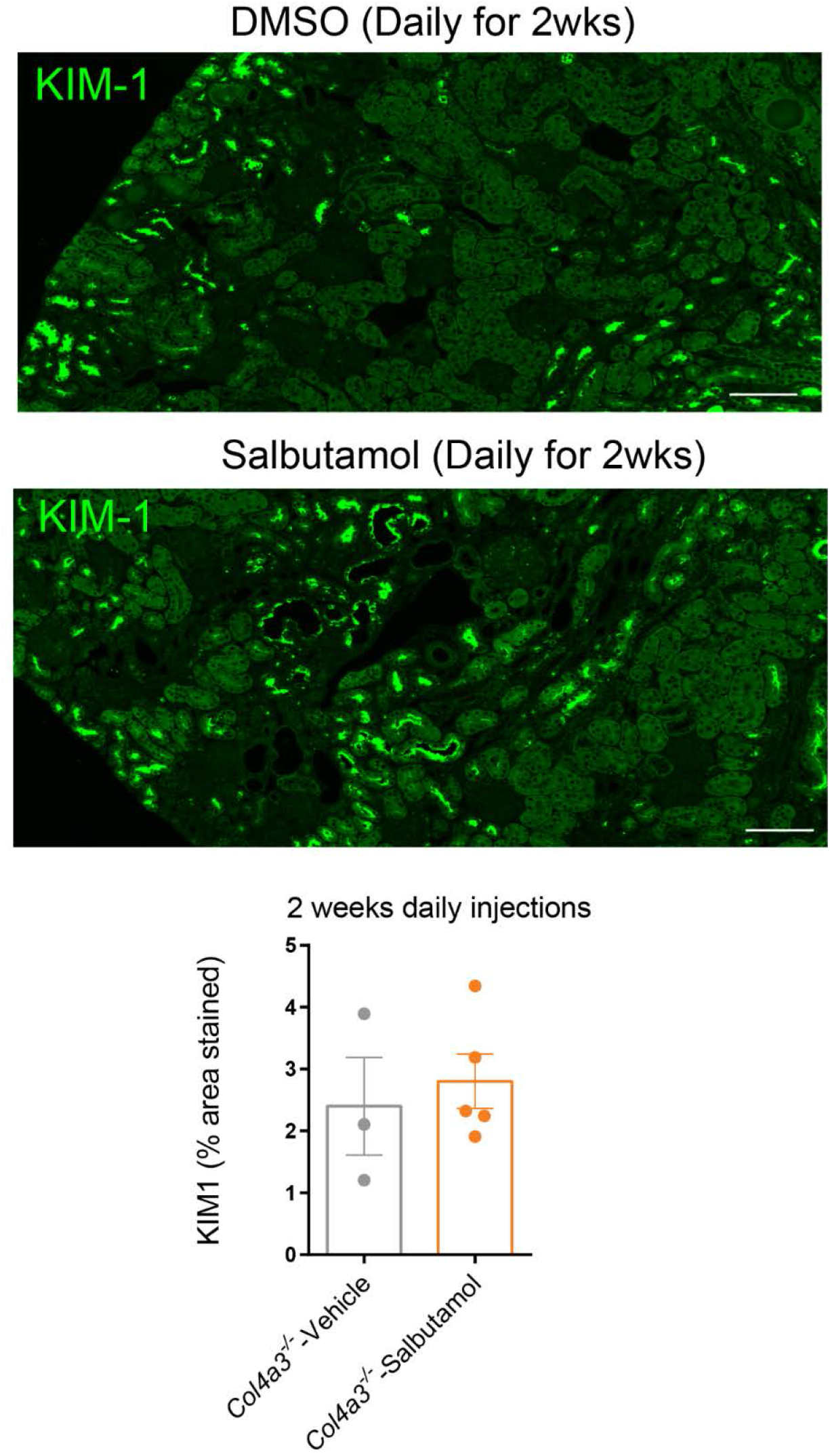
Effects of chronic treatment of Alport *(Col4a3^-/-^)* mice with salbutamol on Kidney Injury Molecule-1 (KIM-1). Quantification of KIM-1 immunostaining showed that chronic administration of salbutamol (100 μg daily dose for 2 weeks) did not change KIM-1 expression levels. Data are mean±SEM. N=3-5 mice per group and 5 images per kidney section. Two independent users quantified the images. No significance was detected using Unpaired Student’s t-test. Images were acquired at 10X using VS120 slide scanner. Scale bar = 100μm.

**Table S1.** Demographic and diagnostic information of the subjects retained for blood collection.

| Code | Gender | Age (years) | Diagnosis and genetics (if known) |
| --- | --- | --- | --- |
| Alport Patient 1 | M | 18.15 | Alport Syndrome, Col4A5 frameshift mutation, stage 5 chronic kidney disease with nephrotic-range proteinuria (highest in the cohort) |
| Alport Patient 2 | M | 17.83 | Alport Syndrome |
| Alport Patient 3 | F | 4.82 | Alport Syndrome, COL4A4 c.3022G>A, p.(Gly1008Arg); steroid-sensitive nephrotic syndrome, no renal insufficiency at the time of collection. |
| Asymptomatic Alport Patient 4 | M | 38.68 | Biopsy-proven clinically asymptomatic Alport with normal renal function. |
| Non-Alport Control 1 | M | 50.15 | Unknown |
| Non-Alport Control 2 | M | 33.92 | Gross hematuria, Kidney stone |
| Non-Alport Control 3 | M | 21.21 | Microscopic hematuria. Congenital Single kidney with hypertension |
| Healthy Control 1 | F | 22.1 | Healthy Individual |
| Healthy Control 2 | F | 22.1 | Healthy Individual |
| Healthy Control 3 | M | 22.9 | Healthy Individual |
| Healthy Control 4 | M | 21.1 | Healthy Individual |

## ONLINE METHODS

### Blood sample collection and induced pluripotent stem cell (iPSC) reprogramming and culture

Blood samples were collected in SepMate tubes from four consenting patients (including three symptomatic and one asymptomatic Alport patients), and 5 healthy control donors and 3 individuals with non-Alport renal disease using established IRB protocols at the University of Miami. A commercial kit was used for peripheral blood mononuclear cells (PBMCs) isolation and culturing (StemPro®-34 SFM Complete Medium, Gibco), together with a kit containing viral vectors that express the Yamanaka factors (Cytotune 2.0, ThermoFisher) for reprogramming PBMCs into induced pluripotent stem cells (iPSCs). Briefly, PBMCs were purified with Lymphoprep density gradient medium and cultured in StemPro®-34 SFM Complete Medium. To generate iPSCs the Cytotune 2.0 Sendai Virus reprogramming kits (Life Technologies) was utilized. Following transduction of PBMCs, the medium was transitioned to MTeSR Plus Medium (StemCell Technologies Cat#05825) until iPSC colonies form in high-attachment 6-well plates coated with Vitronectin (StemCell Technologies Cat#07180). Once iPSC colonies are established, they are maintained and expanded in MTeSR Plus Medium on vitronectin-coated 12-well plates. To establish high-quality iPSC lines, many iPSC colonies (clones) were be expanded and characterized. AT least two clones per sample were validated and used for downstream differentiation experiments.

### iPSC characterization and culturing

iPSCs were plated into Matrigel-coated flasks (Corning, 354277) with mTeSR1 medium (StemCell Technologies, 85850). For passaging, cultures of iPSCs that reached 70-80% confluency were lifted using Gentle Cell Dissociation Reagent (StemCell Technologies, 07174) and transferred to new Matrigel-coated flasks. The iPSCs were validated by flow cytometry and immunocytochemistry using pluripotency markers OCT4, SOX2, NANOG, SSEA4, and TRA-1-60.

### Renal tubular epithelial cell (TEC) differentiation

As previously described by Narayanan *et al*^1^ iPSCs cultured with mTeSR media (Stem Cell Technologies) on Matrigel-coated plates (Corning) were lifted using Gentle Cell Dissociation Reagent (Stem Cell Technologies), pipetted to disrupt the colonies and plated back to Matrigel-coated plates using Renal Epithelial Growth Medium (REGM, Promocell) supplemented with the following supplements: 10 ng/mL BMP2 (StemCell Technologies), 2.5 ng/mL BMP7 (R&D Systems), 10 ng/ml activin-A (StemCell Technologies) and 0.1 μM Retinoic Acid (Sigma). The culture medium was changed every other day for 15-20 days. Then, the morphology of the iPSC-TECs was evaluated and successful differentiation was validated by flow cytometry for tubular markers (KSP and GGT1) and using γ-Glutamyltransferase (GGT) activity assay as described below. HK-2 cells (ATCC, CRL-2190) which are human kidney proximal tubular cells were used as positive control cells for GGT activity assay.

### Morphological and functional characterization of the iPSCs and differentiated TECs

Differentiated TECs were tested for pluripotency markers (SOX-2, SSEA-4, OCT-, TRA-160) as well as tubular epithelial markers (AQP-1, KSP) using immunostaining and flow cytometry to confirm the lack of pluripotency and presence of renal lineage in the differentiated cells. To further validate the generated TECs, γ-glutamyl transferase (GGT) activity, as a functional marker of tubular epithelial cells, was measured in iPSCs and iPS-TECs (at day 20 post-differentiation) from Alport and control subjects using a colorimetric assay kit (MAK089; Sigma-Aldrich) according to the manufacturer’s instructions.

### Flow cytometry

Both iPSCs (prior to differentiation) and iPS-TECs were dissociated with 0.05 trypsin/EDTA (Invitrogen) for 3 minutes at 37 *°*C. The cells were mechanically dissociated by pipetting and/or scraping, filtered through a 70 μm cell strainer (Falcon) then fixed with Cytofix solution (BD Biosciences) for 10 minutes on ice. Cells were then spun down at 1500 rpm at 4C for 5 minutes, washed with Perm/Wash solution (BD Biosciences) and permeabilized using Cytoperm solution (BD Biosciences, 554714) for 15 minutes. Cells were then incubated with conjugated antibodies against pluripotency markers TRA-1-60 (Biolegend 330620, 1:10), SOX-2 (Biolegend, 656112, 1:10), and/or SSEA4 (Biolegend 330418, 1:10) and tubular markers Kidney-Specific Protein (KSP, R&D Systems, NBP2-54567AF488, 1:20) and γ-Glutamyltransferase (GGT1, Biolegend 394304, 1:20) for one hour in a cold room. Corresponding isotype immunoglobulins were used as negative control. Cells were then counted using a flow cytometer (LSR-II; Beckman Coulter) according to the manufacturer’s instructions. The data was analyzed to determine the rate of TRA-1-60-negative, SOX-2-negative cells expressing KSP and GGT. The expression levels of other relevant proteins were also evaluated using flow cytometry as described above including Kidney Injury Molecule (KIM1, Biolegend, 353906, 1:10), and Low Density Lipoprotein Receptor (LDLR, BD Biosciences 565653, 1:100). The data was analyzed using FCS Express software.

### *In vivo* salbutamol and pHrodo-Red LDL cholesterol injections

Alport mice on 129J background were used for daily intraperitoneal injections. A stock solution of 50mM salbutamol (sigma, S8260) was prepared in DMSO, further diluted in 100μL DPBS and was used in acute and chronic studies.

For short term studies, 200 μg of salbutamol as a single injection was used 7-week-old animals with overnight fasting. Animals were euthanized 1h or 3h post-injection. Blood samples were collected for measuring blood urea nitrogen (BUN) and creatinine (CRE) levels, kidneys were collected for histology and protein analysis. Urine samples were collected immediately prior to the injection and at dissection 3h post-injection for albumin and creatinine assays.

For chronic treatment studies, salbutamol was injected daily at 100μg/mouse for 14 days in 6-week-old mice. On the last day of injections, echocardiography and Doppler imaging were performed as described below to assess cardiac function. The next day, the animals were fasted overnight and euthanized, blood samples were collected for measuring BUN and CRE levels, kidneys were collected for histology and protein analysis and urine samples were saved for albumin and creatinine assays.

### *In vivo* salbutamol and pHrodo-Red LDL cholesterol injections

To evaluate the LDL-C uptake *in vivo,* the Alport mice were injected with 100μg of pHrodo Red (pHrodo™ Red-LDL-C) 2 hours prior to dissection. The injection for acute studies was performed 1h after salbutamol administration. Upon euthanasia, the animal was perfused with 10mL of PBS and then kidney tissues were collected in 4% PFA and processed for OCT embedding. Kidney cryosections images were captured at 40X magnification and 0.6X digital zoom using Z-stacking on a Zeiss LSM710 confocal microscope. LDL-C uptake was quantified in renal tubules using NIH’s Image J as the positive red area normalized by total tubular area.

### Echocardiography and strain analysis

Cardiac function was evaluated by a Vevo2100 imaging system (VisualSonics, Toronto, Canada) with a MS400 linear array transducer. All measurements were performed excluding the respiration peaks and obtained in triplicate; the mean value was used for data analysis. The data were analyzed with VevoLab 1.7.1 software (VisualSonics). Briefly, mice were anesthetized with 4% isoflurane at 0.8 L/min flow rate and maintained with 1% isoflurane. Following anesthesia, the mice were fixed in a supine position on a pad with an integrated temperature sensor, a heater, and ECG electrodes. The heart rate was monitored constantly and body temperature was maintained at 37°C during measurement. Depilatory cream was used to remove the fur from the region of interest, and medical ultrasonographic acoustic gel was used as a coupling fluid between the real-time microvisualization scan head and the skin. Image acquisition was initiated with the transducer probe placed along the left sternal border to obtain the parasternal long axis view, which displays both the apex and the outflow tract of the left ventricle. An M-mode gate was placed through the center of the papillary level long-axis view to obtain standard M-mode recordings.

Afterwards, the probe was rotated 90 degrees to obtain and record short-axis images at the level of the mid-papillary muscles. An M-mode gate was placed through the center of the papillary level short-axis view to obtain standard M-mode recordings. Cardiac function parameters examined in B-Mode included: Endocardial Area (d, diastole and s, systole); Epicardial Area (d, s) Endocardial Major (d, s); Endocardial Volume (d, s); Endocardial Stroke Volume; Endocardial %EF; Endocardial %FAC; Endocardial Area Change; Endocardial CO. In M-Mode: IVS, Interventricular septum (d, s); LVAW, left ventricular anterior wall (d, s); LVID, Left ventricular internal diameter (d, s); LVPW, Left ventricular posterior wall (d, s); Left ventricle volume (d, s); CO; %EF; %FS; LV Mass; LV Mass Corrected.

Mitral valve (MV) inflow was assessed from the apical 4 chamber view. The MV measurements performed were the following: MV early wave peak (E), MV atrial wave peak (A), no flow time (NFT), aortic ejection time (AET), isovolumic relaxation time (IVRT), isovolumic contraction time (IVCT), E wave deceleration time (DT). Then Tissue Doppler Imaging (TDI) was performed on mitral valve leaflet tips to measure the velocity of myocardial motion. From TDI we measured the E’ wave corresponding to the motion of the mitral annulus during early diastolic filling of the LV, and the A’ wave which originates from atrial systole during late filling of the left ventricle.^2^ Strain analysis was performed from B-Mode for long and short axes views. The endocardial borders were traced in the end-systolic frame of the 2D images to obtain maximal opposing wall delay, global longitudinal strain (GLS, for long axis), and global circumferential strain (GCS, for short axis).

### Immunostaining for KIM-1

Immunostaining of paraffin-embedded kidney sections was performed for KIM1 (R&D, Cat no. AF1817) at 1:200. The stained slides were then imaged at 20X magnification using an Olympus VS120-L100 Virtual Slide Microscope (Tokyo, Japan), and KIM1 expression was subsequently quantified using Image J.

### Immunostaining for βPix

Immunostaining of paraffin-embedded kidney sections was performed for βPix (SC-136035) at 1:200. The stained slides were then imaged at 20X magnification using an Olympus VS120-L100 Virtual Slide Microscope (Tokyo, Japan), and βPix expression was subsequently quantified using Image J.

### Biochemical assays

Kidney and heart lysates of Alport mice treated with salbutamol or control vehicle were used to measure cAMP levels using a commercial kit (Cat N. ADI-900-163, Enzo Life Sciences). Urine samples were used to measure urinary Albumin to creatinine (Alb/CRE) ratio using a mouse albumin ELISA kit (Cat No. E99-134, Bethyl Laboratories) and a Creatinine Companion ELISA kit (Cat No. 1012, Exocell) according to the manufacturer’s instructions. The levels of blood BUN and CRE were measured in fresh plasma samples by the Comparative Pathology Core Facility at the University of Miami Miller School of Medicine.

### Cell Lines and Transfection

Cell culture media and supplements were obtained from Invitrogen (Carlsbad, CA). Anti-βPix antibodies were from Millipore (Temecula, CA) or Santa Cruz Biotechnologies (Santa Cruz, CA). Antibodies against LDLR, GFP, GAPDH and Myc were purchased from Santa Cruz Biotechnologies (Santa Cruz, CA). Anti-IDOL and rabbit anti-LDLR antibodies were from Abcam (Cambridge, MA). Anti-FLAG antibody was from Sigma-Aldrich (St. Louis, MO). Anti-HA antibody was from BioLegend (San Diego, CA).

HK2, HEK293 and HepG2 cell lines were purchased from American Type Culture Collection (ATCC). HK2 cells were grown in serum-free K-SFM purchased from Thermo Fischer Scientific (Waltham, MA). HepG2 and HEK293 cells were cultured in MEM (Sigma) and DMEM (Invitrogen), supplemented with 10% fetal bovine serum, penicillin (100 units/ml), and streptomycin (100 μg/ml) in a 37 °C humidified incubator with 5% CO2. Transient transfection of cells with mammalian expression vectors was performed using Lipofectamine 2000 (Invitrogen) according to the manufacturer’s instructions.

### Alport Dog Aortic Smooth Muscle Cells (AoSMCs)

Primary canine AoSMCs were isolated from aortic explants of an 8-month-old male with X-linked hereditary nephropathy who had reached end-stage kidney disease (serum creatinine ≥ 5 mg/dL). Cells were cultured for an average of 5 days in 6-well plates pre-coated with fibronectin (18μg/mL), maintained at 37°C (5% CO2, 90% humidity) with DMEM/F-12 culture medias (Gibco, 11960044 and 11765054) supplemented with 20% FBS (Gibco, A3160501), gentamycin (1μg/mL; Gibco, 15750060), sodium pyruvate (1 mmol/mL; Sigma, S8636), and penicillin-streptomycin (100 U/mL; Gibco, 15140122). Canine AoSMCs were expanded into T25 tissue culture flasks when 90% confluency was reached. Media was changed every two days.

### Plasmids

Myc-tagged-β1Pix, β1PixDHm (L238R, L239S), β1PixSH3m (W43K), β1PixΔDH, β1PixΔPH, and β1PixΔ(602-611) plasmids were described previously^3,4^. Lentiviral plasmid for GFP-LDLR and plasmid for FLAG-IDOL were purchased from Genecopoeia. HA-Ubiquitin was obtained from Addgene (plasmid # 18712).

### Immunoprecipitation and Immunoblotting

Cells were transfected with appropriate constructs for 24 h, washed twice in phosphate-buffered saline (PBS), and lysed in a solution containing 20 mM Tris, pH 7.5, 100 mM NaCl, 5 mM MgCl2, 1 mM EDTA, 1% Triton X-100, 1 mM sodium fluoride, 1 mM sodium vanadate, 1 mM phenylmethylsulfonyl fluoride, 1 μg/ml pepstatin, and 1 μg/ml leupeptin. Equal amounts of proteins were separated using either 8% or 4-12% sodium dodecyl sulfate-polyacrylamide gel electrophoresis, transferred onto nitrocellulose membranes (Millipore), immunoblotted with appropriate antibodies, and visualized using enhanced chemiluminescence (SuperSignal West Femto; Amersham Thermo Scientific). For immunoprecipitation, anti-LDLR antibody was added to cell lysates for 2 h, followed by addition of protein G-agarose beads for an additional hour. Beads were washed three times in lysis buffer. Immunoprecipitated proteins were released from beads by boiling in 1× sample buffer for 5 min and analyzed by immunoblotting. Total cell lysates (TCL) were analyzed to assess over-expression of constructs. Expression of Myc-β1Pix mutants was verified by immunoblotting with anti-Myc antibody.

### Cell treatments

Differentiated iPS-TECs, HK2 cells and were seeded in appropriate plates and treated with short-acting β_2_AR agonist salbutamol (Sigma, S8260) or an equivalent amount of DMSO as control vehicle, and then used for functional or biochemical assays as described below.

### LDL cholesterol uptake assay

LDL cholesterol uptake assay was performed on HK2 cells, AoSMCs from XLHN dog and the Alport and control subjects’ iPS-TECs to examine the effects of β_2_AR agonism. As detailed in our previous work,^5^ iPS-TECs were plated on a 24 well plate at 30,000 cells/well using the supplemented REGM media. HK2 cells and XLHN dog AoSMCs were plated at 10000 cells/well on 24-well plates. The following day (for HK2 cells and AoSMCs) and on differentiation day 19 for iPS-TECs, the cells were lipoprotein-starved by changing the media to the respective media without FBS. After 24 hours of starvation, cells were treated with salbutamol or vehicle as described above and placed in the incubator for 30 minutes. Then, each well was immediately treated with 10μg/mL pHrodo Red-Labelled LDL cholesterol (Thermofisher Scientific, L34356) and the plate was placed in the Live Cell Imaging System (Incucyte, Essen BioScience) for live imaging (16 images/well hourly for 10-20 intervals) using phase and red channels. The data was analyzed using IncuCyte Zoom software to obtain area normalized total red object intensity as we previously described.^5^

